# Anisotropic Subcutaneous Response During Fingertip Normal and Tangential Loadings

**DOI:** 10.1101/2023.04.12.536577

**Authors:** Guillaume H.C. Duprez, Benoit P. Delhaye, Delannay Laurent

## Abstract

The subcutaneous mechanical response of the fingertip is highly anisotropic due to the presence of a network of collagen fibers linking the outer skin layer to the bone. Yet, the impact of this anisotropy on the fingerpad deformation had not been studied until now. This issue is here tackled using a two-dimensional finite element model of a transverse section of the finger. Different hypotheses about the orientation of the fibers are considered: radial (physiologic), circumferential, and random (isotropic behavior). The three variants of the model are assessed using experimental observations of a finger leaning on a flat surface. Predictions relying on the physiological orientation of fibers align best with reality. Moreover, the orientation of fibers plays a crucial role in determining the distribution of internal strain and stress. These factors, combined with the abrupt shift in contact pressure during the transition from sticking to slipping, represent important sensory cues for partial slip detection. This is valuable information for the development of haptic devices.

## I. Introduction

HEN grasping objects or exploring surfaces, our perception of touch is ensured by biological sensors, called *tactile mechanoreceptive afferents*, which densely innervate the fingertips [1], [2]. The related neuronal activity may be probed under a large variety of stimuli. However, in such experiments (see for review [1], [3]–[5]), it remains unclear which information exactly is communicated to the brain. One of the important challenges in the interpretation of mechano-transduction is that afferents are embedded at some depth (from *∼* 500*μ*m to several mm) inside the finger pad and they hence witness subcutaneous strains which are likely to differ from those observed along the skin outer surface [6].

Various modeling approaches have aimed to simulate the deformation of a finger leaning on a flat surface. Analytical solutions have been derived for the simplest loading scenarios, by considering that the finger is either a frictionless elastic half-space (Cerruti problem) [7]–[9], or an elastic membrane filled with an incompressible fluid (waterbed model), which better predicts surface deflection under line load [10]. Somewhat more realistic simulations were achieved using the finite element method. The response of the soft biological tissues was then most often considered linear elastic (assumption of infinitesimal strains), sometimes accounting for the multilayered structure of the skin [11], [12] or the topology of the fingerprint [13]. The entire hand was modeled in [14]. Only a few studies [15], [16] assumed a hyperelastic material behavior, which suits better physiological tissues under finite isochoric deformation. Most of the previous studies involved monotonic loadings but a few of them did consider the fingertip response under oscillatory loads [17]–[19].

All of the latter studies assumed an isotropic material response, disregarding the fact that the network of collagen fibers, which is hosted by subcutaneous tissues, is likely to induce significant anisotropy. Indeed histological observations have evidenced that collagen fibers are not randomly distributed. They rather are “radially” aligned: one fiber end is anchored on the bone and the other end comes close to reaching the skin [20]. If the deformation of the fingertip is such that some collagen fibers are stretched, these fibers are expected to sustain high tensile stresses along their axis [21]. The famous HGO hyperelastic model [22] accounts for the asymmetry of the fiber response under, respectively, compressive and tensile axial loads. It also predicts the anisotropic macroscopic response resulting from the preferential alignment of the collagen fiber network, and it fulfills volume preservation as expected in soft biological tissues [23], [24].

The present study aims to assess to which extent the anisotropy of subcutaneous tissues is likely to influence the deformation of a finger leaning on a flat surface as well as the build-up of internal stresses and the pressure distribution along the surface of contact. The finite element model described in Section II is used for the simulation of laboratory experiments in Section III. The discussion of results in Section IV precedes the conclusion.

## II. Description of the numerical model

The simulations involve a finger leaning against a horizontal plate which is considered perfectly flat and infinitely stiff. This is a fair assumption as the plate used in experiments (Fig. 1(a-b)) is made of glass. Fig. 1(c) defines the characteristic dimensions of the finger ellipsöidal cross-section. Contact with the plate is established by prescribing a vertical displacement (along ***ê***_*y*_) to the bone, assuming that only the surrounding subcutaneous tissue can deform. Friction is simulated by displacing the bone along the in-plane horizontal direction ***ê***_*x*_ while maintaining a fixed amplitude of the vertical contact force. The Coulomb coefficient for friction is assumed constant and its value is determined (in the next section) based on experimental measurements of the ratio between the tangential (horizontal) and normal (vertical) contact forces.

**Fig. 1.**
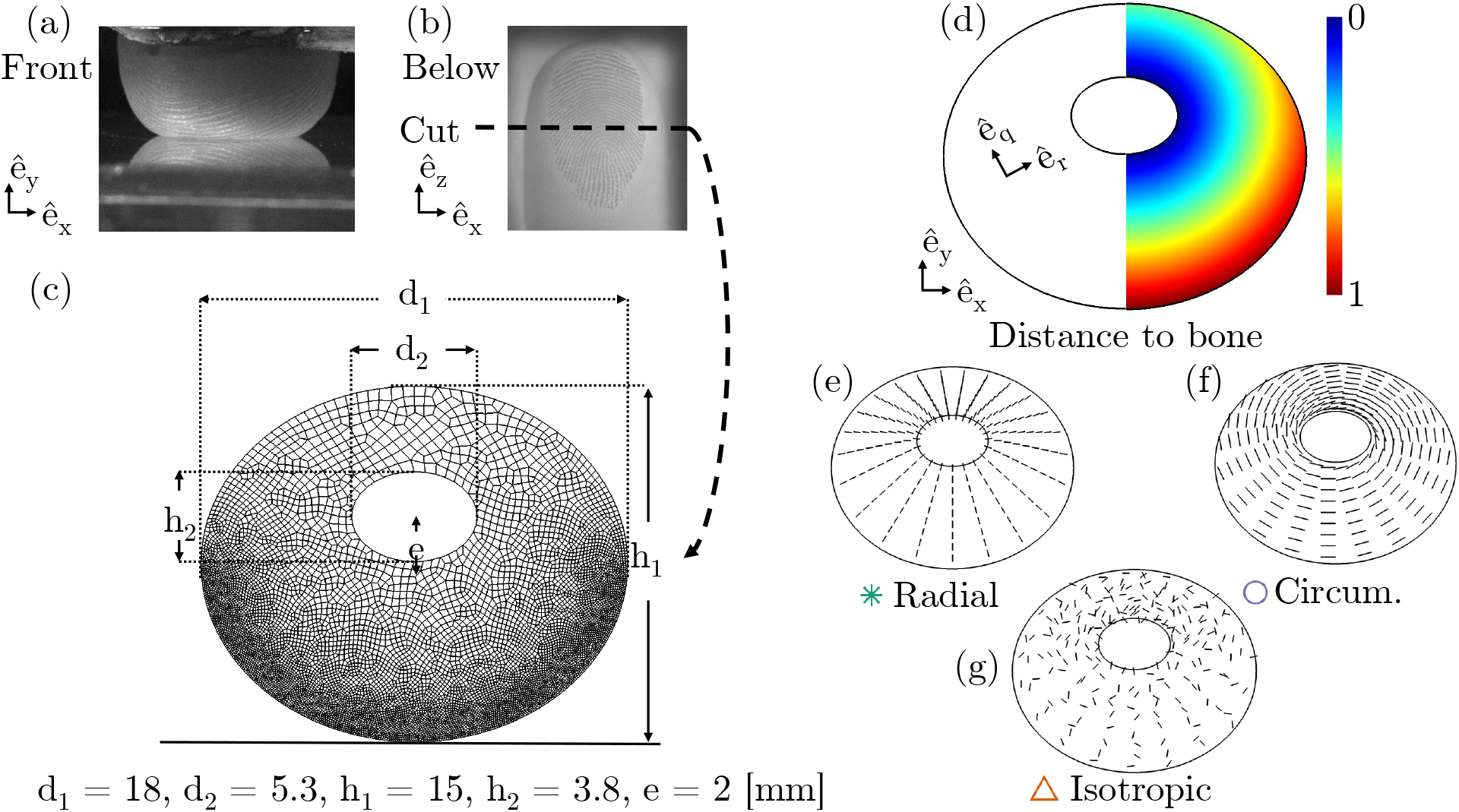
Modeling of a 2D fingertip reinforced by fibers network. (a-b) Fingertip compression against a glass plate seen from the front or from below; (c) 2D model of the fingertip in contact with a rigid line with geometrical dimensions; (d) Gradient of the distance to the bone; (e) Radially oriented fibers; (f) Circumferentially oriented fibers; (g) Randomly oriented fibers ensuring isotropic macroscopic behavior.

Simulations are carried out using the Abaqus finite element (FE) code **abaqus**. The FE mesh is generated using the gmsh suite [25] and the element size is refined in the region which comes in contact with the stiff plate (Fig. 1(c)). Generalized plane-strain conditions are adopted, which means that deformation is uniform along the out-of-plane direction ***ê***_*z*_. Simulations are reproduced twice using different values of the out-of-plane stiffness. Hybrid, 4-noded elements (CPEG4H in abaqus) are used to overcome the numerical (locking) issues which may arise when a material undergoes finite strains under constant volume. This implies that the hydrostatic stress is computed at every location of the mesh as a Lagrange multiplier enforcing that the divergence of the displacement field be nil (according to the weak formulation of the mechanical equilibrium equations, which is typical of FE solutions).

Inside the subcutaneous tissue, a significant increase of stiffness is expected when collagen fibers are stretched in the course of the loading. The latter is accounted for by relying on the HGO anisotropic hyperelastic model [22]. If we consider that, at any location across the FE mesh, collagen fibers are precisely aligned along *N* directions (no local dispersion relative to these directions), the hyperelastic strain-energy density function *W* of the HGO model may be expressed as follows:

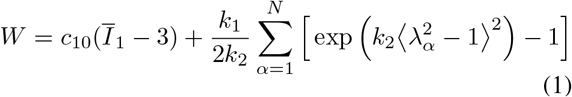

Here conventional denominations are used: the parameters *c*_10_ and *k*_1_ are stiffnesses representative of, respectively, the isotropic ground matrix of the subcutaneous tissue and the collagen fibers, whereas the *k*_2_ parameter is dimensionless. The scalar variable 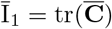 is the first invariant of the isochoric part of the right Cauchy-Green strain tensor: 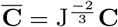 where **C** = **F**^*T*^ **F** and **F** is the deformation gradient tensor. The scalar variable 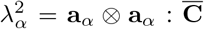 corresponds to the square of the stretch of the collagen fibers that are aligned with the unit vector **a**_*α*_. As collagen fibers do not sustain compressive axial loads, the argument of the exponential is under Macaulay brackets: ⟨*x −* 1 ⟩ = *x −* 1 if *x >* 1, and ⟨*x −* 1⟩ = 0 otherwise.

In order to investigate the influence of the anisotropy of subcutaneous tissue on the deformation of the finger cross-section, three modeling hypotheses are tested regarding the orientations of collagen fibers. They are illustrated in Fig. 1(e-g). The first situation is closest to the histological observations [20]: collagen fibers follow the shortest path from the bone to the skin, which is referred to as a *radial* network of fibers. The second situation treated is a fictitious (unrealistic) one in which collagen fibers are aligned *circumferentially* around the bone. In the third set of simulations, the distribution of fibers is random so that the average response is isotropic. In practice, the definition of appropriate fiber orientation(s) at every integration point of the FE mesh proceeds as follows:

- The “radial” network considers a single fiber orientation (*N* = 1) which corresponds to the shortest path to the inner ellipse (i.e. to the bone). The distance of each node of the mesh to the bone is thus computed and the unit “radial” vector (***ê***_*r*_ in Fig. 1 (d-e)) is computed based on the gradient of such distance field.
- The “circumferential” fiber orientations are defined inside the same cross-section, using the unit vector ***ê***_*θ*_ which is perpendicular to ***ê***_*r*_ as shown in Fig. 1(d,f). Here again a single local orientation of collagen fibers is used at every integration point (*N* = 1).
- The isotropic response is obtained by considering three fiber orientations (*N* = 3) at every integration point. The first fiber orientation is selected randomly in the 2D section (Fig. 1(g)) and the other two orientations are tilted +120° and *−*120° relative to the first one. Such a definition of fiber orientations ensures isotropy and reduces noise.

## III. Results

The two-dimensional FE model is assessed based on a comparison of the numerical predictions to experimental observations of the deformation of a fingertip in contact with a glass plate (Fig. 2). To start, the parameters of the model are determined by ensuring that all three variants of the model reproduce the evolution of the contact force as a function of the finger displacement when the loading is normal to the plate, i.e. along ***ê***_*y*_ in Fig. 1. The reference experiments, labelled *A* and *B* in Fig. 2, were carried out in [26]. Only one parameter (*k*_1_) is adapted when switching from one variant of the model to another. This parameter is proportional to the volume fraction of collagen fibers. Experimental trends are properly reproduced when *k*_1_ = 150[kPa] in case of the radial fibers network, and when *k*_1_ = 3[kPa] if the model relies on either the random or the circumferential fibers network. The other parameters are assigned values that do not depend on the orientation of collagen fibers: *c*_10_ = 3[kPa] represents the stiffness of the background matrix and *k*_2_ = 15[-] determines the rate of increase of the stiffness when collagen fibers are stretched. These parameter values are all within the usual range when the HGO model is used to represent soft biological tissues [22]–[24].

**Fig. 2.**
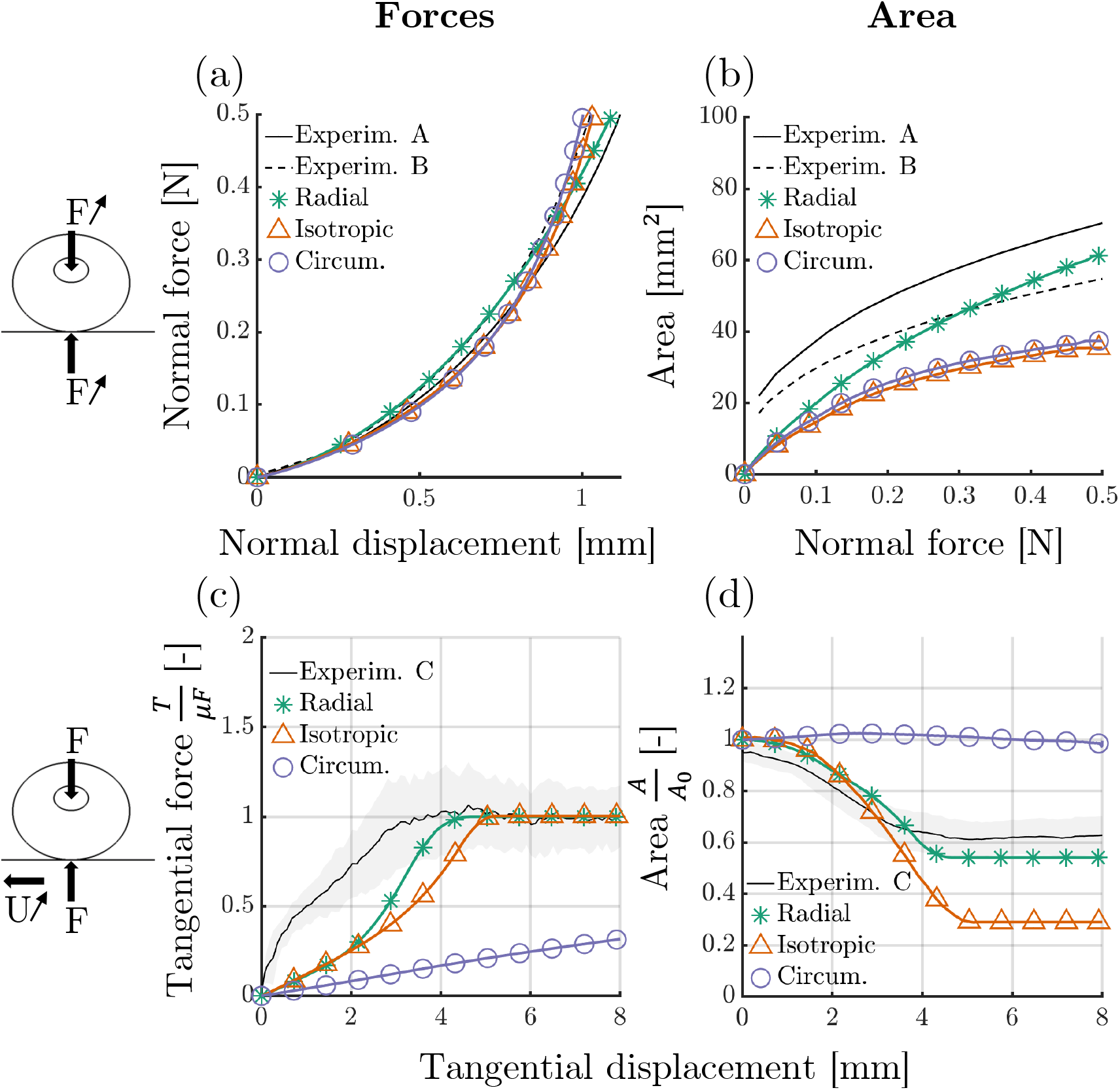
Comparison of models predictions with normal (Experim. A-B, measured at 30*°*and 45*°*fingertip inclination angles [26] and tangential loading data (Experim. C, [27]). (a) Normal force-displacement; (b) Gross contact area during normal loading; (c) Normalized tangential forces trend during tangential loading; (d) Relative contact area during tangential loading; The light grey area in (c) and (d) represents the standard deviation of the experimental data.

As Fig. 2(b) shows, still in the case of normal loading, the three variants of the model lead to different predictions of the increase of the contact area as a function of the contact force. The simulations being performed in 2D, the contact area *A* is here deduced from the predicted contact length *L* while assuming a circular contact area: 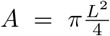. Among the three predictions, the experiment trend is best captured when the model assumes a radial network of collagen fibers.

The three variants of the model are now evaluated by simulating friction under tangential loading. The normal component of the contact force is kept fixed at 0.5 [N]. The friction coefficient is *μ* = 1, which conforms to experimental measurements [28]. According to both the model and the experiment, both the tangential component of the force and the contact area evolve as function of the horizontal displacement of the bone. Both the force and the area reach a constant value when fingertip sliding is initiated. In the experiments performed in [27], the finger moved along either the ulnar or the radial direction (both under 0.5 [N] normal force). The average of the two measurements is used here in order to assess model predictions. The material parameters of the HGO model are identical to those used in the first set of simulations (which considered normal loading). In Fig. 2(c) the experimental and predicted tangential forces are normalized with respect to their respective sliding threshold. Fig. 2(d) depicts the corresponding evolution of the contact area. Among the three variants of the model, the worst predictions (those differing most from the experiment) are produced when collagen fibers are circumferentially aligned. The variant of the model with radial fibers is the one performing best, in particular with regard to the predicted contact area under sliding.

Figure 3 shows the profiles of the pressure and the shear stress which are predicted along the line of contact at the glass-fingertip interface. The solid lines correspond to the predictions after normal loading, whereas the profiles represented with a dashed line are predicted after a tangential displacement U = 2.5[mm]. After normal loading (when U is nil), the variant of the model with a radial fiber network predicts a contact pressure with a flattened bell profile, whereas the circumferential and isotropic networks of collagen fibers lead to Hertzian profiles with higher amplitudes. After a tangential displacement U = 2.5[mm] (dashed lines), the pressure profiles become asymmetric and the peak amplitude is shifted towards the trailing edge of the fingertip. The transition is most sudden in the case of the radial fiber network and much more subtle in case of circumferentially oriented fibers. Similar observations are made when considering the profiles of contact shear stress. According to Fig. 2(c) and (d), at this amplitude of lateral displacement, partial slip is initiated only if the collagen fibers have either radial or random orientations.

**Fig. 3.**
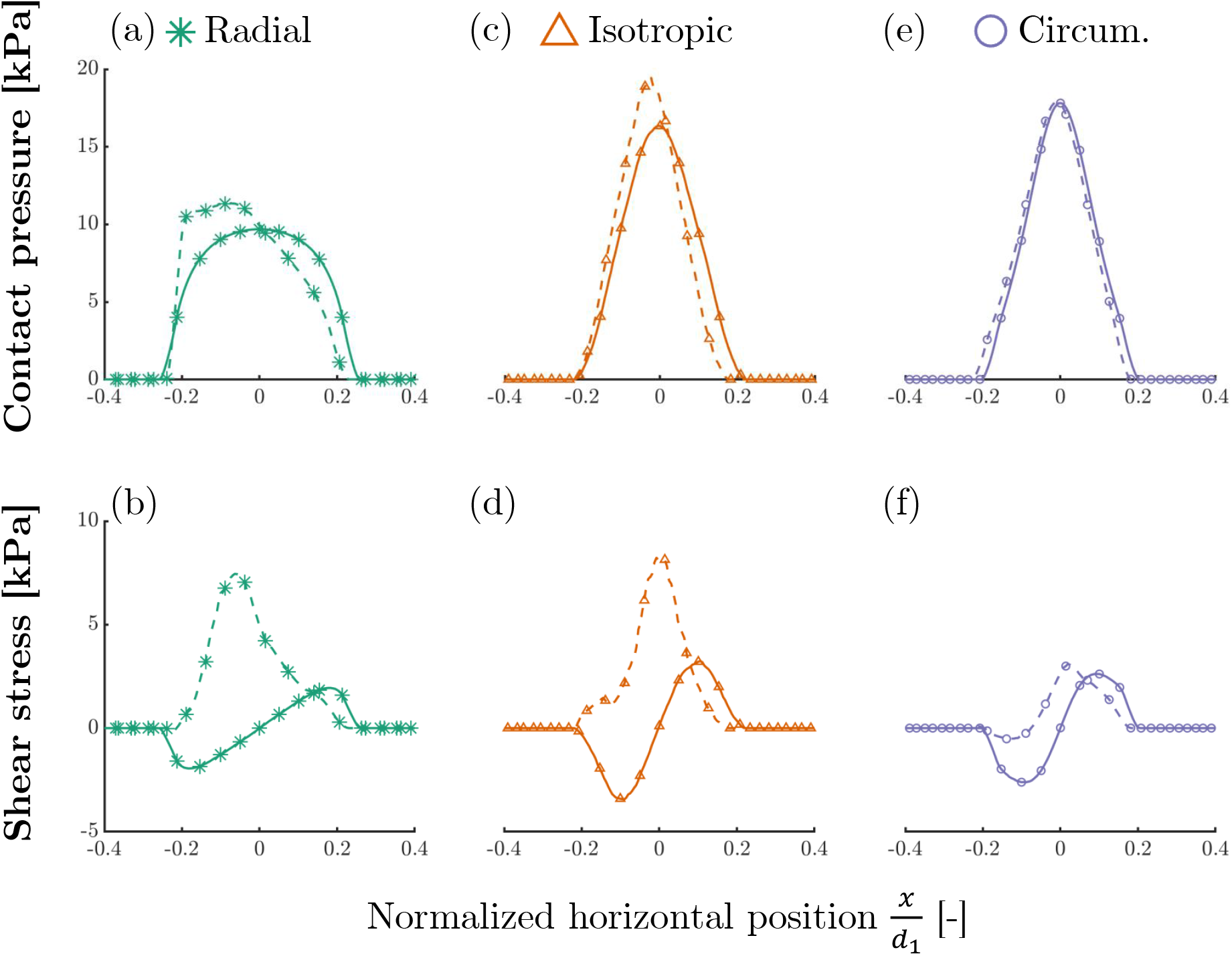
Evolution of the pressure and shear stress along the contact line between fingertip and rigid surface. The pressure profile is symmetric and the shear stress distribution is skew-symmetric under pure normal loading (continuous lines). When contact involves both normal pressure and tangential displacement U = 2.5 [mm] (dashed lines), the maximum pressure is shifted to the trailing side of the finger. Symbols represent a fraction of the extracted data points value.

Figure 4 shows how the anisotropy of the mechanical response influences the heterogeneous deformation of subcutaneous tissues. The deformation fields are analyzed by mapping two variables *λ*_*r*_ and *λ*_*θ*_, which correspond to the local material stretch along, respectively, the radial and circumferential directions. Values larger and lower than 1 correspond, respectively to elongation and contraction along the given direction. In these subfigures, which compare the different variants of the model, we may notice that stretches are impeded (or at least hindered) in the direction of collagen fibers. For instance, the connection of the soft tissue to the bone is stiffer in the case of the radial network of fibers (the values of *λ*_*r*_ around the bone are lowest with this variant of the model).

**Fig. 4.**
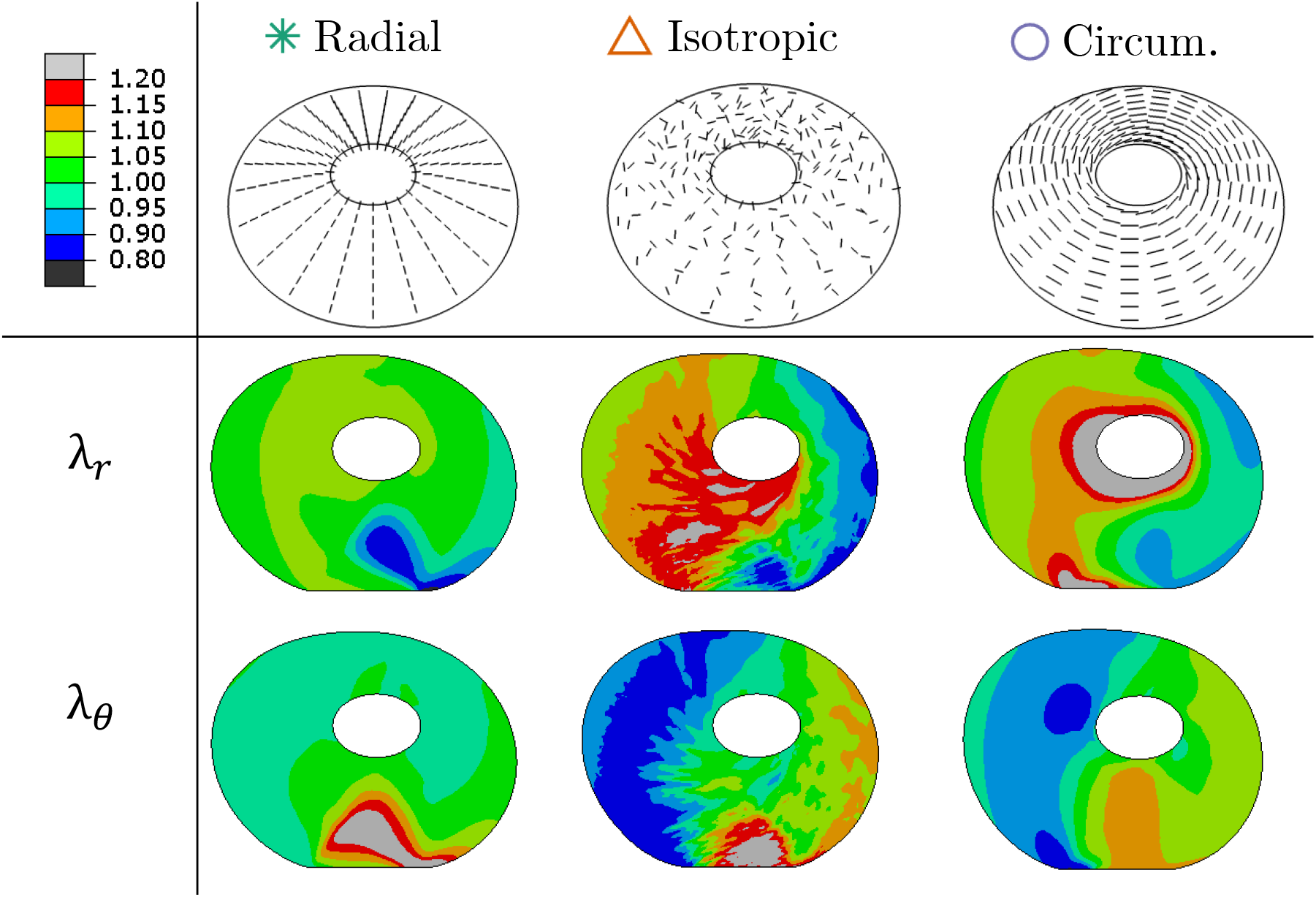
Local elongation or contraction of the material along the radial (*λ*_*r*_) and circumferential (*λ*_*θ*_) directions. In the three simulations (involving different networks of collagen fibers), the contact of the finger with the plate involves a normal displacement of 1 [mm] and a tangential displacement of 8 [mm].

## IV. Discussion

The development of haptic devices requires a thorough understanding of the physiological processes likely to explain the sense of touch. The present study demonstrates that, when a fingertip leans on a surface, the anisotropy of subcutaneous tissues greatly influences the strain and stress fields in the neighborhood of the tactile afferents that trigger neuronal activity. According to the model, the local stretches experienced by such biological sensors vary greatly, depending on the orientations of the network of collagen fibers (Fig. 4). Radial fiber networks, as observed in histological observations, also induce a sudden change of the contact pressure and shear stress (Fig. 3) early in the transition from sticking to slipping. This likely has major impact on tactile feedback. Indeed, tactile afferents are considered highly sensitive to the local pressure and to its rate of change [3], [29]. The occurrence of an asymmetric pressure profile before full slip might be a crucial feedback in order to maintain a stable grasp during manipulation [6], [30]–[33].

The model proposed here relies on some highly simplifying assumptions but its predictions seem realistic. Experimental measurements of the load displacement curve and the evolution of the contact area of a fingerpad leaning on a rigid surface are properly reproduced, while relying on model parameter values in the same range as those identifies in many other studies using the HGO model in order to compute the response of soft biological tissues. The numerical predictions are closest to experimental trends when the orientation of collagen fibers corresponds to histological observations (Fig. 2). As collagen fibers sustain much better axial elongation (tensile stresses) than compression, the physiological fiber configuration, i.e. the radial network, lowers the macroscopic stiffness under compression along the main direction of loading (normal to the rigid surface). This accelerates the growth of the contact area, flattening the profile of contact pressure. The latter has so far not been studied in great detail experimentally, but the available data show similar trends [1].

The numerical simulations are here performed in 2D, while assuming uniform strain in the out-of-plane direction, i.e. generalized plane strain. It had been observed that the amplitude of the out-of-plane deformation does not influence the main trends in the numerical results. The 2D assumption seems justified given the in-plane distribution of collagen fiber orientations and the fact that the finger in contact with the plate moves along a direction parallel to the cross-section that is modeled (Fig. 1). The simulation of more general loading scenarios would require adapting the model from 2D to 3D. Experimental campaigns justifying such 3D modeling are still under progress [6], [34].

Another potential improvement of the model would be to account for the heterogeneity of subcutaneous biological tissues [16]. In the present study, the material response is considered homogeneous apart from the local orientation of collagen fibers. In reality, not only the orientation, but also the density of collagen fibers varies from place to place. The fiber density is highest under the nail and this is likely to impede the “rolling” phenomenon, i.e. the circumferential motion of the skin relative to the bone induced by friction with the plate under tangential loading. The present model also disregards the layered structure of the skin itself. The outermost viable epidermis and dermis are stiffer than the subcutaneous tissues, thus forming a thin “membrane” which is likely to have some influence on the local stress and strain fields. The biological mechanoreceptors are located at the junction between the viable epidermis and the dermis. More precisely, Merkel cells (associated with SA-I afferents) are found at the tip of inner ridges (mirroring the fingerprint ridges) and Meissner corpuscles (associated with FA-I afferents) are nested inside the dermal papillae [35], [36]. Such arrangement surely pursues a goal and it motivates the development of a microscale modeling complementing the present one.

## V. Conclusion

The mechanical anisotropy of subcutaneous tissues has a major influence on both the distribution of contact stresses and the overall deformation of a fingerpad under normal and tangential loading. This is a direct consequence of the nonlinear and directional stiffening of stretched collagen fibers. In the present study, the predictions of a two-dimensional FE model are compared to experimental observations of a fingerpad leaning on a rigid surface, leading to the following conclusions:

- As compared to two alternative modeling hypotheses, the laboratory experiments are reproduced most accurately using the model variant which considers physiological orientations of collagen fibers (radial network).
- The pressure profile along the line of contact of the fingerpad with the surface has a realistic bell shape when collagen fibers are oriented radially and this pressure profile changes suddenly in the transition from sticking to slipping. The latter is likely to produce efficient feedback when grasping objects.
- The mechanical anisotropy of subcutaneous tissues influences also the deformation in the close neighborhood of the tactile afferents triggering neuronal activity. This should be accounted for in the development of haptic devices.

## Aknowledgements

LD and BD are mandated by the Belgian National Fund for Scientific Research (FSR-FNRS).

